# Slower prefrontal metastable dynamics during deliberation predicts error trials in a distance discrimination task

**DOI:** 10.1101/2021.02.01.429249

**Authors:** Danilo Benozzo, Giancarlo La Camera, Aldo Genovesio

## Abstract

Previous studies have established the involvement of prefrontal cortex (PFC) neurons in decision processes in many task contexts. Single neurons and populations of neurons have been found to represent stimuli, actions, and internal deliberations. However, it is much less clear which underlying computations are affected during errors. Neural activity during errors can help to disambiguate confounds and clarify which computations are essential during a specific task. Here, we used a hidden Markov model (HMM) to perform a trial-by-trial analysis of ensembles of simultaneously recorded neurons from the dorsolateral prefrontal (PFdl) cortex of two rhesus monkeys performing a distance discrimination task. The HMM segments the neural activity into sequences of metastable states, allowing to link neural ensemble dynamics with task and behavioral features in the absence of external triggers. We report a precise relationship between the modulation of the metastable dynamics and task features. Specifically, we found that errors were made more often when the metastable dynamics slowed down, while trial difficulty influenced the latency of state transitions at a pivotal point during the trial. Both these phenomena occurred during the decision interval and not following the action, with errors occurring in both easy and difficult trials. Thus, modulations of metastable dynamics reflected a state of internal deliberation rather than actions taken or, in the case of error trials, objective trial difficulty. Our results show that temporal modulations of PFdl activity are key determinants of internal deliberations, providing further support for the emerging role of metastable cortical dynamics in mediating complex cognitive functions and behavior.

## Introduction

The role of prefrontal cortex in decision making has been demonstrated using a variety of tasks (Kim and Shadlen, 1999; Hoshi et al., 2000; Freedman et al., 2001; Tanji and Hoshi, 2001; Genovesio et al., 2012; Maoz et al., 2013; Marcos and Genovesio, 2016; Rich et al., 2018), including tasks with decisions in the temporal (Genovesio et al., 2009) and spatial (Genovesio et al., 2011) domains. In particular, we have shown previously that prefrontal neurons encode the decision about which stimulus was presented farther from the center in a distance discrimination task (Genovesio et al., 2011), as well as the transformation of goals into action (Marcos et al., 2019). Prefrontal neurons also code for the duration and distance magnitude to be maintained in working memory using domain-specific representations of distance and duration in PFdl (Marcos et al., 2017). However, in the spatial discrimination task the effect of trial difficulty has not been addressed and previous analyses could not unmask which type of failure could account for the response errors. It is possible that such information could be found in the collective activity of ensembles of neurons. For example, ensembles of monkey premotor neurons transition more quickly to a new state in easier trials during a vibro-tactile discrimination task (Ponce-Alvarez et al., 2012). One might also suspect that ensemble activity is related to behavioral performance and not just trial difficulty. Little is known, however, about the substrate of behavioral errors in terms of collective activity of ensembles of neurons. Examining collective neural activity during errors might also help to understand what generates the switch in coding from correct to erroneous decisions observed at the level of single neurons (Genovesio et al., 2011). A particularly promising approach to investigate the relationship between decision processes and ensemble neural activity is by using a hidden Markov model (HMM) to segment the neural activity into sequences of discrete, metastable states. In its more frequent applications to neural data so far (Abeles et al., 1995; Jones et al., 2007; Kemere et al., 2008; Ponce Alvarez et al., 2008; Ponce-Alvarez et al., 2012; Mazzucato et al., 2015; Engel et al., 2016; Sadacca et al., 2016; Linderman et al., 2016), the HMM assumes that the neural dynamics proceeds as a sequence of hidden states, wherein each state is a collection of firing rates across simultaneously recorded neurons. Reliable sequences of HMM states have been found in frontal (Abeles et al., 1995), gustatory (Jones et al., 2007; Mazzucato et al., 2015), premotor and motor areas (Ponce-Alvarez et al., 2012) of primates and rodents in the context of different tasks. State sequences also seem to underlie internal states of attention (Engel et al., 2016), expectation (Mazzucato et al., 2019), and deliberation during decisions (Rich and Wallis, 2016). In this work, we investigate the nature of neural ensemble dynamics in the PFdl of monkeys performing the distance discrimination task of (Genovesio et al., 2011, 2012). In this task the monkeys were required to choose which of two stimuli sequentially presented was farther from the center of a computer screen, Fig. 1. We focused on two main questions: whether the neural activity in PFdl could be characterized as metastable and, if so, what are the links among metastable state durations, state transition times, trial difficulty, and task performance (correct vs. incorrect trials). We found that PFdl activity can be characterized as a sequence of metastable states, some of which coding for the relative distance of the two stimuli from the center, based on stimulus features or order of presentation. Most notably, however, we found that the mean duration of metastable states was longer before errors. Thus, a slowdown of the metastable dynamics after the presentation of the second stimulus was typically associated with an incorrect response. We also found a link between trial difficulty and hidden state transitions in correct trials (Ponce-Alvarez et al., 2012). Specifically, the first state transition after the second stimulus was faster in easier trials, even though the reaction times were not different. This suggests that transition times reflect the level of trial difficulty even when the delay before a response is too long to affect the reaction time. Overall, these results show a potential role of metastable dynamics in PFdl during a decision process in the spatial domain, adding to accumulating evidence for a role of metastable dynamics in neural coding and cognition (La Camera et al., 2019).

**Figure 1:**
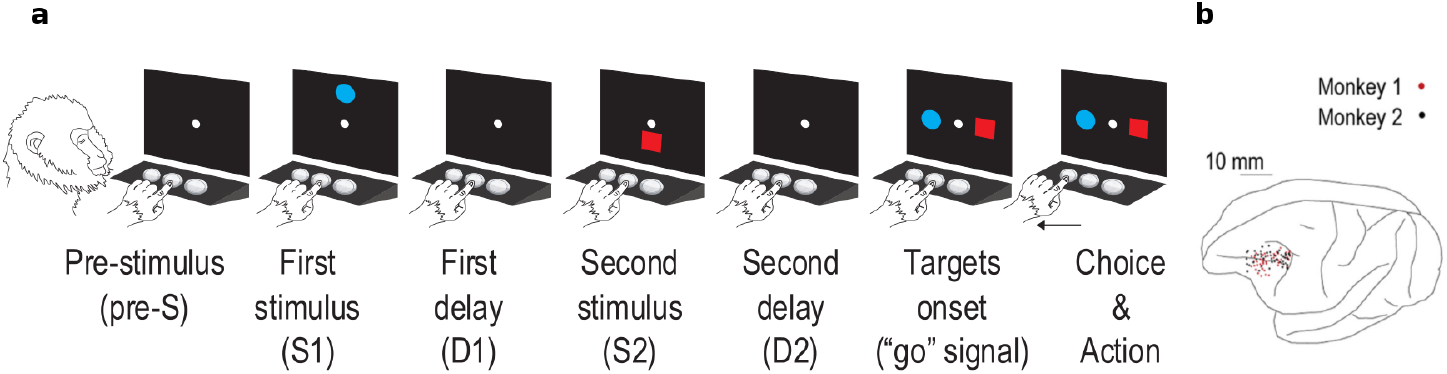
Experimental paradigm and neural recordings sites. (a) Sequence of task events within a trial: each trial started when the monkey touched the central switch, leading to the appearance of the central stimulus (reference point) which lasted for 400 or 800 ms (pre-S). After this pre-stimulus period the first stimulus (S1) was presented. S1 was followed by a variable delay D1 of 400 or 800 ms which lasted until the second stimulus (S2) appeared. S2 was followed by a second delay (D2) of 0, 400, or 800 ms. Both S1 and S2 were presented for 1000 ms and placed either above or below the reference point at 8 to 48 mm (8 mm step) from the reference point. After both stimuli reappeared (placed horizontally), and the monkey had to touch the switch below the stimulus that appeared farther from the reference point (the blue circle in the example trial in the figure). Correct responses were rewarded with 0.1 ml fluid while errors were followed by an acoustic feedback. The stimulus feature (blue circle/red square), position (above/below the reference point), distance, and target position (left/right) were pseudo-randomly selected. (b) Penetration sites. Composite from both monkeys, relative to sulcal landmarks. See Methods for details.

## Results

### Metastability of prefrontal neural activity during distance discrimination

In order to find neural correlates of behavior in the ensemble dynamics, we first characterized the dynamics by performing an HMM analysis of the ensemble activity of correct and error trials (see Fig. 2 and Methods for details of the procedure). We found that ensemble neural activity was well described by sequences of metastable states, where each state was defined as a collection of firing rates across simultaneously recorded neurons, with the number of states varying between 2 and 5 across sessions. As in previous accounts (Abeles et al., 1995; Jones et al., 2007; Mazzucato et al., 2015, 2019; Ponce-Alvarez et al., 2012) state transitions varied trial-by-trial and were not necessarily locked to external relevant events. This can be illustrated by warping time so that events in different trials occur simultaneously (Recanatesi et al., 2020), while onset and offset times of states are variable, as shown in Fig. 3a for an illustrative session (see Methods for details). From this plot, it is apparent that the blue state tends to occur after S2 and GO, while the yellow state tends to occur after the decision (RT). However, the time at which transitions occur varies from trial to trial, as documented in the mentioned studies. Moreover, during trial epochs in which no clear external events occur, such as halfway after S2 and during the delay period, state transitions are much more variable, possibly reflecting the internal dynamics of state deliberation (more on this later).

**Figure 2:**
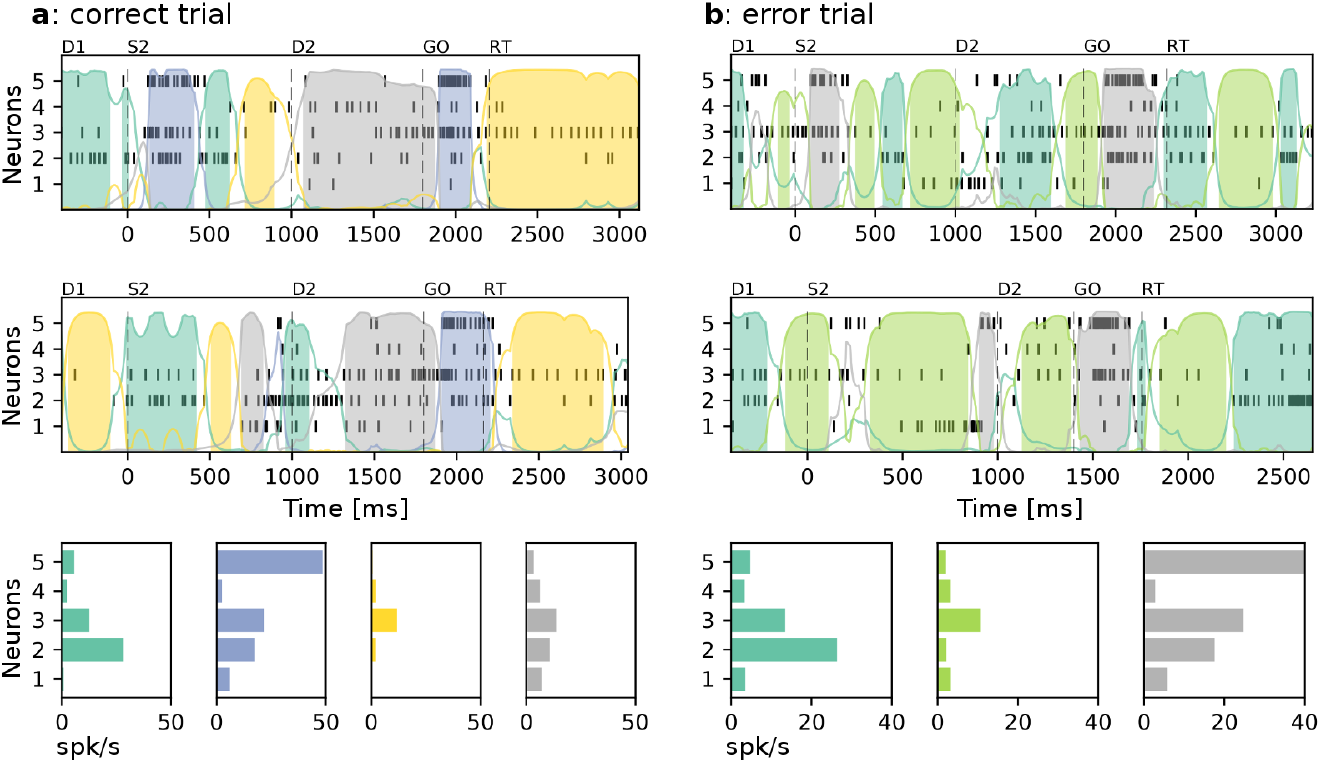
Representative trials of a neural ensemble of simultaneously recorded cells. Each raster plot shows the spiking activity of each recorded neuron from 400 ms before S2 presentation until the beginning of the following trial. Colored curves represent the posterior state probabilities and the assigned states are indicated with colored areas. Insets (bottom panels) show firing rate vectors associated with each state (same color code as in corresponding top panels). (a) Correct trials. (b) Error trials. Note that similar colors in (a) and (b) do not correspond to similar states since the HMMs were performed independently on correct and error trials (see Methods). Key: D1: first delay (after S1); S2: second stimulus; D2: second delay; GO: targets appear on screen; RT: reaction time.

**Figure 3:**
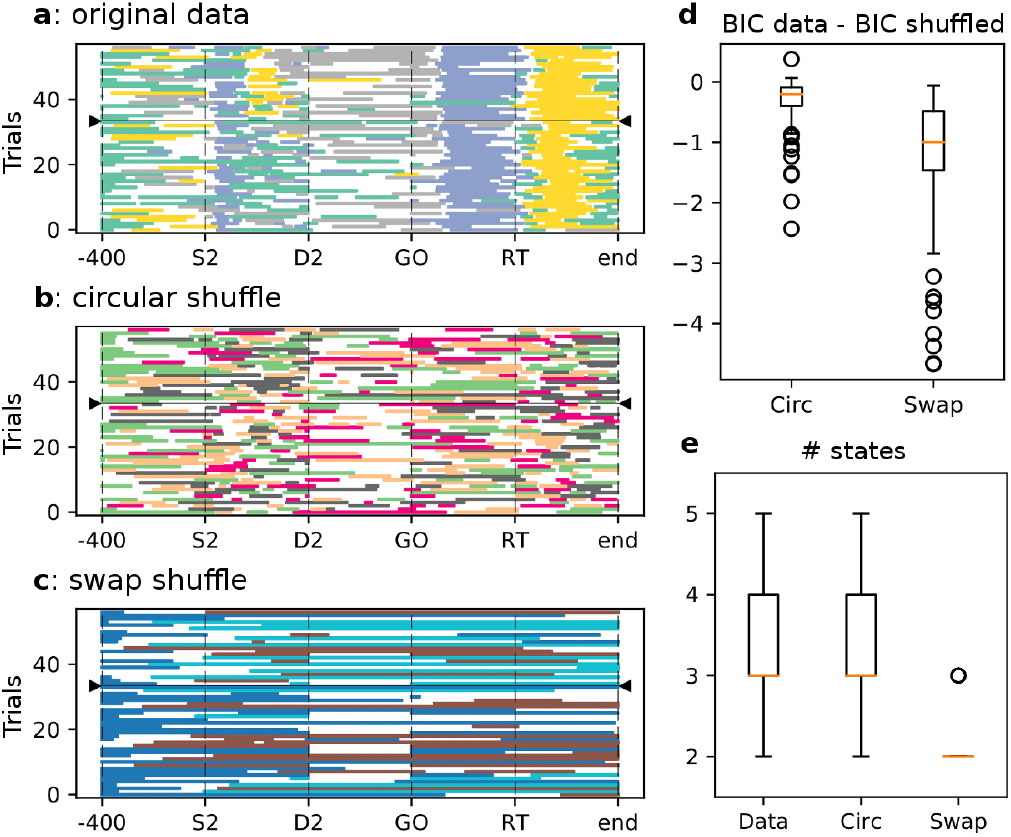
Comparison of HMM state sequences for original and shuffled datasets. (a,b,c) Comparison of HMM state sequences for original and shuffled datasets in one representative session (only trials with S2 in the UP position shown; similar plots are obtained in other conditions). Trials were grouped according to the relative distance based on stimulus features (blue circle vs. red square), and the separation between the two groups is indicated by the black horizontal line with borders marked by triangles (with red trials above the line). The time in each trial has been warped (stretched or shrunk, see Methods) so as to align the 4 events S2, D2, GO, RT. (a) HMM model of original data reveals (i) the presence of a coding state for relative distance based on stimulus features (the yellow state appearing between S2 and D2 only in trials above the group separation line, coding for “red square farther”) and (ii) reliable state transitions at relevant event times. (b) HMM model of circularly shuffled data shown in panel a. As a consequence, state sequences appear scrambled and the coding states are lost. (c) HMM model of swap-shuffled data shown in panel a. Compared to panel a, sequences are not orderly despite the presence of fewer states. (d) Boxplots of the difference in BIC score between the fits to the original data and shuffled data across sessions (*p* < 0.001, Wilcoxon signed-rank test). A smaller score indicates better fit. (e) Optimal number of inferred states across sessions for original data (left), circular-shuffled data (middle) and swap-shuffled data (right). Note that similar color in panels a-c does not imply the same state.

The orderly sequences observed in Fig. 3a were not an artifact of the HMM analysis, as they disappear in shuffled datasets. We used two types of shuffling procedures introduced in (Maboudi et al., 2018), which we call circular and swap shuffles (see Methods). As shown in Fig. 3b-c, both shuffling procedures disrupted the temporal alignment between state sequence and task events. The difference with the original data is especially evident following S2, GO and RT. Moreover, model selection had a larger score (smaller BIC) in the original data than in the shuffled datasets (Fig. 3d), revealing worse HMM fits when the data were shuffled. The analysis also shows a significant reduction of the optimal number of states after swap-shuffling the data (between 2 and 3 compared to 2-5 with a median of 3 for the original data; Fig. 3e). The comparison with shuffled datasets shows that the presence of metastable activity and the potential meaning of hidden states is not an artifact of the HMM analysis.

To investigate how the hidden states relate to relevant task variables, we searched for “coding states”. We define a coding state as a hidden state that tends to recur significantly more often in a specific task condition compared to other task conditions (Mazzucato et al., 2019). To illustrate this notion, in Fig. 3a-c trials were grouped according to the features (blue circle vs. red square) of the farther stimulus from the center. The separation between the two groups is indicated by the black horizontal line with extremities marked by triangles. Note that, during S2 and prior to D2, the yellow state appears only in trials that are above the horizontal separation line, and therefore it is a coding state for relative distance based on stimulus features (i.e., “red farther” or “blue closer”) in the current trial (*p* < 0.05, *χ*^2^ test; see Methods for details). We found coding states for the features of the farther stimulus in 22% of the sessions (two example sessions are shown in Fig. 4a). This is a significantly larger fraction than the 6% found in the circularly shuffled datasets (*p* = 0.0006, *χ*^2^-test; not enough state transitions occurred in the swap shuffled data). We also found evidence for coding states for relative distance based on the order of presentation (“S2 farther or closer from the center”) in 19% of sessions (two example sessions shown in Fig. 4b). For comparison, only 5% of sessions in shuffled datasets had coding states for relative distance based on the order of presentation, a significant difference (*p* = 0.002, *χ*^2^-test). We did not find coding states significantly associated with target position (left vs. right) between GO and RT (6% of sessions vs. 2% of shuffled sessions, *p* = 0.089, *χ*^2^-test). This set of comparisons with shuffled datasets shows that the presence of metastable activity, as well as the nature of hidden states as coding for relevant task variables, are not an artifact of the HMM analysis.

**Figure 4:**
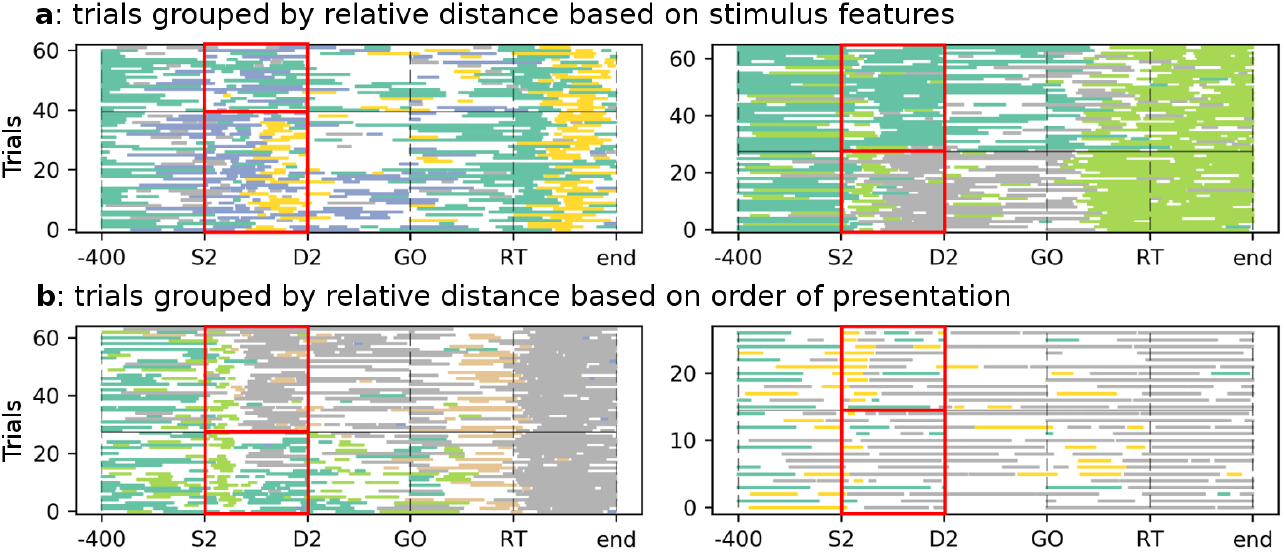
Examples of sessions with significant coding states. (a) Coding states for relative distance based on stimulus features (blue circle vs. red square) during S2 (red box) in correct trials for 2 example sessions. Coding states are the dark green and yellow states in the left panel and the dark green and grey states in the right panel. (b) Coding states for relative distance based on order of presentation during S2 (S2 farther vs. S2 closer) in correct trials for 2 example sessions. Coding states are the dark green, orange and grey states in the left panel and the yellow state in the right panel. In both panels, trials were grouped according to the coded variable (as in Fig. 3a) and highlighted by the red box. The same colors in different panels do not imply the same state.

### Mean state durations predict incorrect trials

As shown in Fig. 5a, reaction times were longer in error trials than in correct trials. However, reaction times were neither correlated with task difficulty (Jonckheere’s trend test, *p* = 0.10) nor with the time of first transition after S2 (Spearman’s rank correlation *r* = 0.05, *p* = 0.31; see the next section for details). Based on recent findings on the role of metastable dynamics in cognitive processes (Mazzucato et al., 2019), we considered the possibility that longer reaction times in error trials may result from a global modulation of the neural dynamics. For example, a slowdown of the dynamics would manifest itself through longer state durations and, therefore, less frequent transitions among hidden states. To test this hypothesis, we compared the mean durations of the hidden states in both conditions (correct vs. error) over a time interval going from 400 ms prior to S2 onset until the end of the trial (Fig. 5b). In analyzing state durations, only states occurring after S2 were considered. Based on the criteria reported in Methods, we analyzed 56 sessions. On average, there were 82 ± 35 correct trials and 22 ± 11 incorrect trials per session (mean ± SD). The optimal number of states of the HMM in each session ranged between 2 and 5 across sessions, with a median of 3 for both correct and incorrect trials. We found that the mean state duration after S2 was significantly greater in incorrect trials, 374.0 ± 9.4 ms (mean±sem), than in correct trials, 321.8 ± 3.7 ms (*p* < 0.001, two-sided Mann-Whitney U test; Fig. 5b).

**Figure 5:**
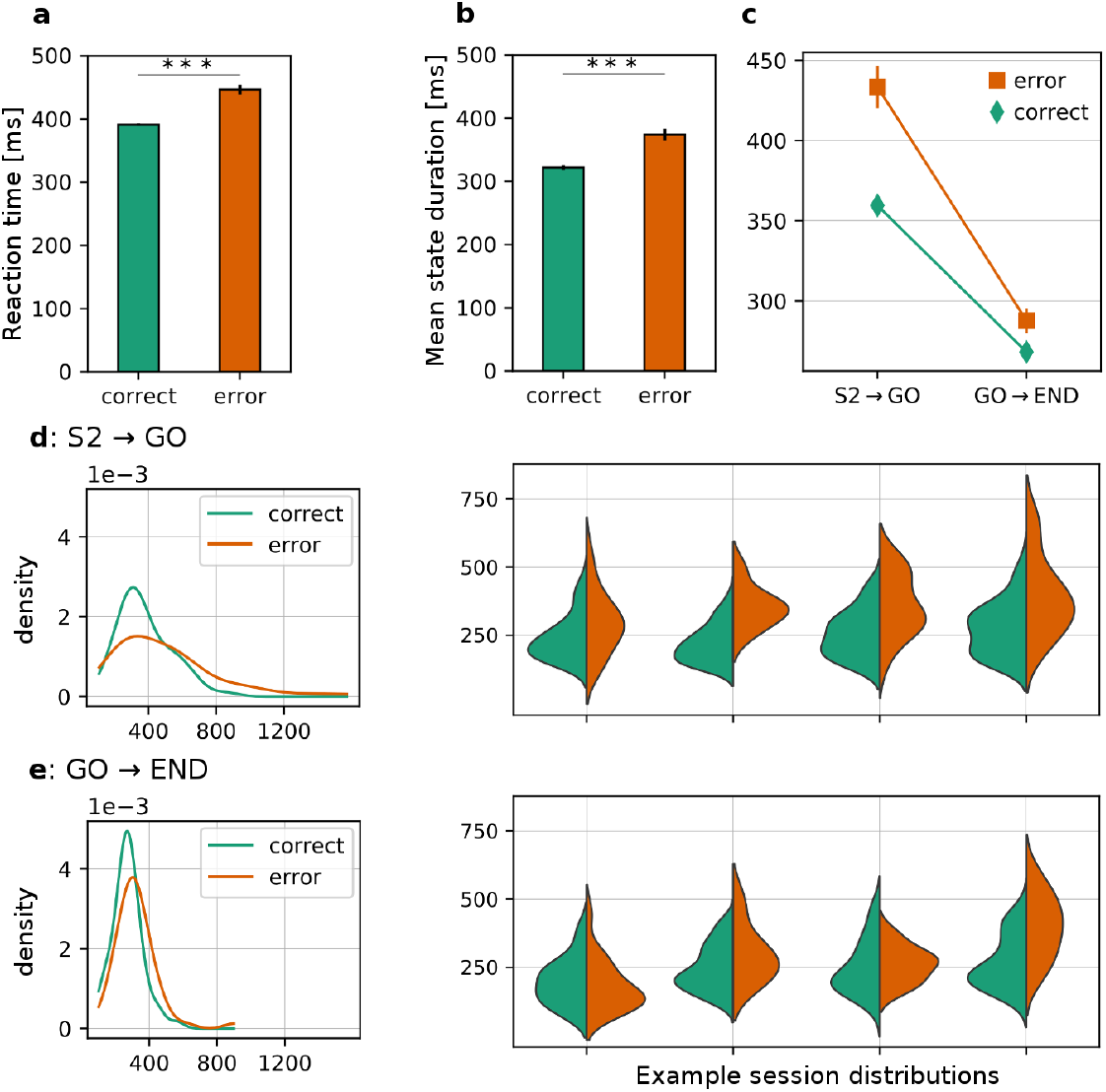
Comparison of reaction times (a) and mean state durations (b) between correct (green) and incorrect (red) trials. The mean value±sem is reported. *** = *p* < 0.001, two-sided Mann-Whitney U test. (c) Comparison of mean state durations in the S2-GO vs. the GO-END intervals, divided in correct and error trials. The plot shows the interaction plot of the two-way ANOVA with factors “trial type” (*p* < 10^−12^, *F* (1) = 50.9) and “temporal window” (*p* = 0, *F* (1) = 363.6, *p*(interaction) < 10^−5^, *F* (1) = 16.3). (d) Left panel: distributions of mean state durations across sessions in the S2-GO period in correct and error trials. Right panel: distributions of state durations in 4 example sessions (green: correct trials; red: error trials). (e) Same as panel (d) for the GO-END period.

A longer state duration in incorrect trials is in line with the hypothesis that the slower reaction times in incorrect trials are associated with a global slowdown of the dynamics. However, we found only a negligible correlation between mean state durations after GO and reaction times (Spearman’s rank correlation *r* = 0.05 for correct trials, *p* = 0.0009, and *r* = −0.01 for error trials, *p* = 0.75). It is possible that cortical slowdown occurs during the deliberation period only, and it directly affects the decision but not the reaction time (which occurs after the GO signal).

To test this hypothesis, we computed the mean state durations in two periods, from S2 to GO (deliberation period) and from GO to the end of the trial (included the intertrial interval), and performed a 2-way ANOVA with trial type (correct/incorrect) and temporal window as factors (Fig. 5c). Both the main effects of trial type and temporal window were significant (*p* < 0.001): overall, state durations were longer during the S2-GO period compared to the GO-END period (in both correct and incorrect trials; see panels d-e for the distributions of mean state durations). However, the reduction in mean state duration observed in the GO-END period (compared to the S2-GO period) was larger in error trials compared to correct trials (Fig. 5c; *p*(interaction) < 10^−5^, *F* (1) = 16.3; Fig. 5d-e shows distributions of mean state durations in the two time periods, together with examples from single sessions). This result shows that slowing down of cortical dynamics occurred mostly in the S2-GO period, i.e., in the period when internal deliberation occurs (360 ms in correct trials vs. 440 ms in error trials). After the GO, state durations dropped below 300 ms in both correct and incorrect trials, suggesting that cortical slowdown occurred during deliberation, rather than during the motor action (accordingly, state durations after GO were not correlated with trial difficulty, i.e., with |S2-S1|: Jonckheere’s trend test gave *p* = 0.06 for error trials and *p* = 0.21 for correct trials).

We validated this result by repeating the HMM analysis with a balanced number of correct and error trials within each session. Since error trials were always fewer than correct trials, a random subsample of correct trials was selected in each session, and the procedure was repeated 20 times. During the deliberation period (S2-GO), we found longer state durations in error trials in 19 out of 20 subsampling repetitions (*p* < 0.001, Wilcoxon signed-rank test). As a control, we also performed the same analysis after shuffling the data (circular and swap shuffle, see Methods and Fig. 3). When the data were circularly shuffled, the mean state duration in error trials was found to be longer than in correct trials in about half of the shuffled dataset (11 out of 20) while it was found to be shorter in the other half (*p* = 0.39, Wilcoxon signed-rank test), consistent with the null hypothesis of no difference in state durations between correct and error trials. The ratio 19/20 is significantly larger than the ratio 11/20 obtained in the shuffled data (*p* = 0.0035, *χ*^2^ test for proportions), suggesting that our result is a true property of the data. In the swap-shuffled data we mostly found 2 states (Fig. 3e), which resulted in only a handful of trials with at least 2 transitions in each session, precluding a meaningful analysis of the state durations in this case.

To further quantify the link between mean state durations and performance, we decoded the monkey performance in a fictitious 2AFC task in which we are presented with a correct and an error trial, and for each such pair, we predict that the error trial is the one with longer mean state duration (see Methods for details). This analysis was performed in those 35 out of 56 sessions with significantly longer state durations in the S2-GO period during error trials, and it correctly predicted the performance in (72 ± 16)% of the trials (area under the ROC, mean ± SD). A similar analysis using reaction times rather than state durations gave similar results, (72 ± 12)%. On a trial-by-trial basis, state durations or reaction times would predict the behavior of the monkey in 72% of the trials, and although this is not remarkable in terms of predictive performance, it is well above chance level (*p* < 0.001 in a 2AFC with *n* ≥ 46 trials, binomial test).

### Neural correlates of trial difficulty

After examining the relationship between neural dynamics and performance, we examined the effect of trial difficulty on metastable dynamics. Despite some non-significant trend before GO (not shown), average state duration was not correlated with trial difficulty (between S2 and GO: *p* = 0.059, Jonckheere’s trend test; linear regression slope=−1.25, *p* = 0.24; after GO: *p* = 0.28, Jonckheere’s trend test; linear regression slope= −0.22, *p* = 0.70). In search for a more significant neural correlate of trial difficulty, we followed (Ponce-Alvarez et al., 2012) and looked at the first transition to a new state after S2 onset (fTaS2). The hypothesis was that, regardless of mean state durations, transitions after S2 onset would occur sooner in easy trials than in difficult trials. Fig. 6 shows two examples of HMM analysis for one difficult (Fig. 6a) and one easy (Fig. 6c) trial. The HMM was first performed with only 2 states and centered on a 1400 ms window around the second stimulus S2, specifically, from 400 ms before to 1000 ms after S2 offset, and during correct trials only. To increase statistical power, the HMM analysis was performed separately on trials with S2 appearing above the central stimulus and on trials with S2 appearing below the central stimulus (however, an HMM with all trials lumped together gave similar results; see below). Sessions were then selected based on the ability of the HMM to decode the monkeys’ decision (see Methods). Only sessions with a significant decoding performance were kept for further analysis: 41 out of 61 sessions with S2 appearing above the central stimulus and 34/61 sessions with S2 appearing below, comprising on average 27.6±12.8 trials per session (mean ± SD). Decoding performance was significantly better than in shuffled datasets, where the same analysis was performed after randomly shuffling the class labels (S1>S2 vs. S2>S1, see Methods). In this case, the decoding performance was significant only in 7 out of 61 (S2 above, *p* < 3 · 10^−10^, *χ*^2^ test) and in 6/61 sessions (S2 below, *p* < 7 · 10^−8^, *χ*^2^ test). As hypothesized, the first transition time after S2 correlated with trial difficulty, with faster transitions occurring in easier trials (Fig. 6e). Specifically, we divided all trials in 5 groups according to the difference in spatial distance between S1 and S2, and found a significant decreasing trend of fTaS2 with increasing |S2-S1|(*p* = 0.006, Jonckheere’s trend test (Bewick et al., 2004)). Slope of linear regression fit = −1.74 (*p* = 0.015, two-sided Wald test with t-distribution; an HMM with S2 appearing both above and below the central stimulus gave similar results: linear regression slope = −1.058, *p* = 0.015). This trend disappeared when the data were shuffled (not shown; *p* = 0.28 for circular shuffle and *p* = 0.46 for swap shuffle, respectively; Jonckheere’s trend test).

**Figure 6:**
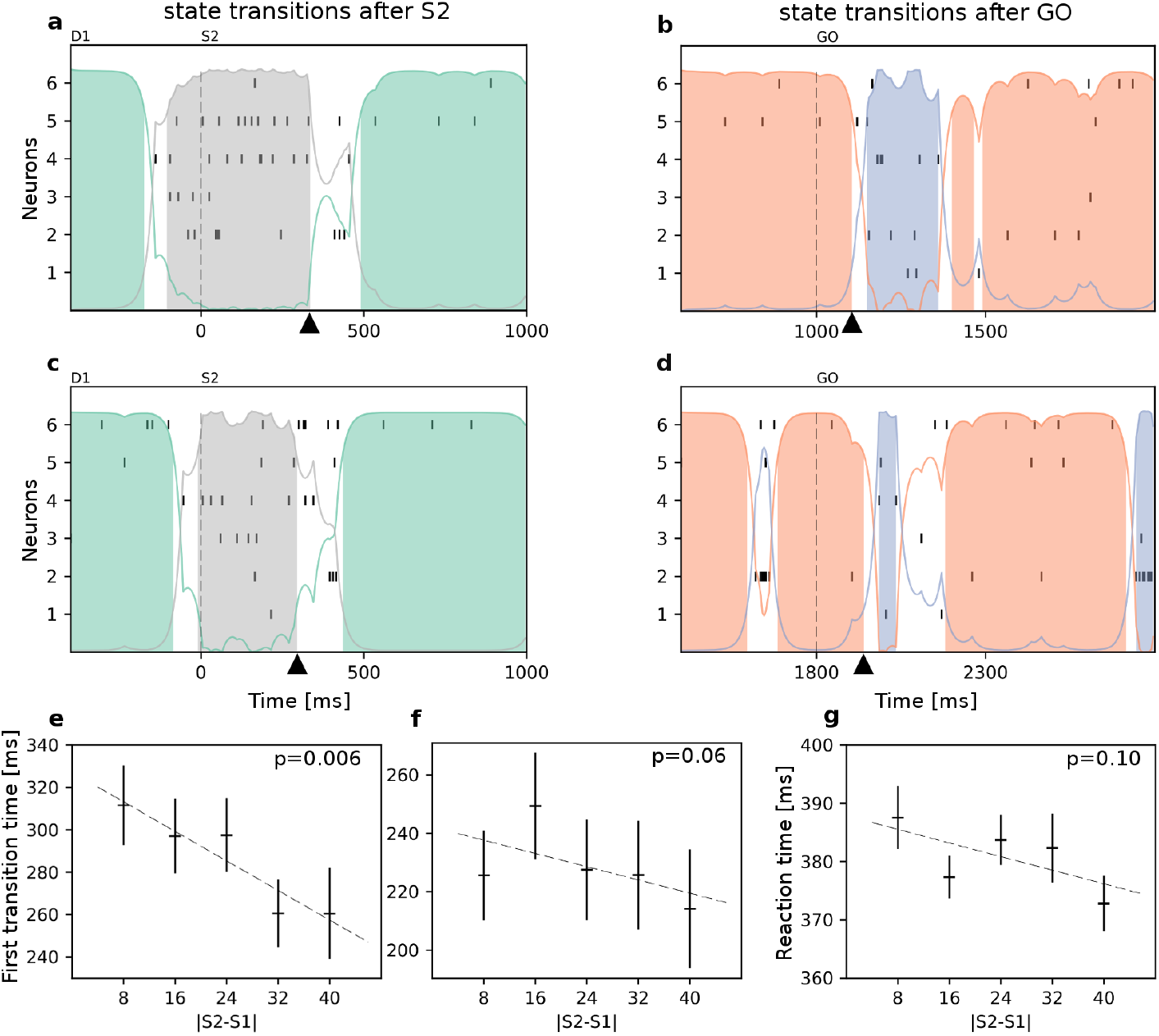
Analysis of first transition time after S2 (fTaS2). (a) Example trial with rasters and HMM segmentation of a neural ensemble of 6 cells recorded simultaneously for one difficult trial with |S2-S1|=16. Each raster plot shows the spiking activity of each recorded neuron from 400 ms before S2 until the end of the stimulus presentation. Same conventions as in Fig. 1. The triangle on the horizontal axis marks the first transition time after S2 (fTaS2). Vertical dashed line: S2 onset. (b) Same trial as in (a) analyzed with an HMM around the GO signal. A time window of 1400 ms was used, 400 ms before and 1000 ms after GO cue. (c) Same as in (a) for an example of easy trial with |S2-S1|=40. (d) Same as in (c) with HMM analysis around the GO signal. (e) fTaS2 vs.|S2-S1|plot (mean±sem) shows a significant trend of fTaS2 with trial difficulty (*p* = 0.006, Jonckheere’s trend test). (f) First transition time (mean±sem) after the GO signal vs. |S2-S1|. No significant trend test, *p* = 0.06. (g) Reaction times (mean±sem) vs. trial difficulty, |S2-S1|. No significant trend test, *p* = 0.10.

These results mirror previous results in motor and dorsal premotor cortex of monkeys (Ponce-Alvarez et al., 2012) and suggest that the signature of a longer process of deliberation in PFdl is a longer transition out of the state present at S2 onset. Since spatial target selection and actions occur after GO, when presumably the decision process has already occurred (except, perhaps, in some of the very difficult trials), faster transition times after S2 would not be expected necessarily to correlate with faster reaction times (the time from GO to action). Indeed, we found that fTaS2s and reaction times were uncorrelated (Spearman’s rank correlation *r* = 0.05, *p* = 0.31). In keeping with this interpretation, we performed a similar analysis for the transition time after GO (Fig.6b-d), and found no relationship with trial difficulty (*p* = 0.06, Jonckheere’s trend test; Fig. 6f). In this case, the HMM analysis was performed in a time interval of −400 ms and +1000 ms around the GO signal. Similarly, the reaction times (time between GO and action) did not depend on trial difficulty (*p* = 0.10, Jonckheere’s trend test; Fig. 6g).

## Discussion

Metastable dynamics of cortical circuits is emerging as a flexible framework to interpret an increasing number of neural phenomena and related behaviors including working memory (Abeles et al., 1995; Ponce-Alvarez et al., 2012), selective attention (Engel et al., 2016) and decision tasks (Rich and Wallis, 2016) in monkeys; navigation (Maboudi et al., 2018), expectation (Mazzucato et al., 2019) and decision tasks (Miller and Katz, 2010) in rodents; and even task related information in human decision making (Taghia et al., 2018) (see La Camera et al., 2019, for a recent review).

Here, we found that the neural activity of ensembles of PFdl neurons of monkeys performing a distance discrimination task are well described as sequences of metastable states. Among those states, some specifically code for the relative distance of the two stimuli from the center, based on stimulus features or order of presentation. Most importantly, this study has uncovered a new role of discrete-state metastable dynamics in the PFdl of monkeys performing a distance discrimination task. Our main results suggest that (i) incorrect decisions correspond to a slow-down of the metastable dynamics preceding the action, regardless of trial difficulty; and (ii) a slower state transition after S2 occurs in difficult correct trials, compared to a faster transition in easy correct trials. We begin by discussing the latter.

### Effect of task difficulty on state transitions

Single units in the PFdl contain information about the decision on which stimulus is farther from the central stimulus (Genovesio et al., 2011). It is not clear if they also encode trial difficulty. Based on previous studies in primary motor and premotor cortex (Pardo-Vazquez et al., 2008; Ponce-Alvarez et al., 2012) and prefrontal cortex (Kim and Shadlen, 1999), we expected that PFdl neurons activity would be affected by trial difficulty. We followed Ponce-Alvarez et al. (2012) and looked for this information in the ensemble dynamics of PFdl neurons as modeled by a HMM. We found that in difficult trials the state transition following the second stimulus, but not the GO signal, had a later onset compared to easier trials. Thus, the state transition latency reflected trial difficulty only in the deliberation period and not later during the transformation from goal to action. This may also explain why, although slower in error trials, reaction times were not correlated with trial difficulty.

Our results are analogous to those found by Ponce-Alvarez et al. (2012) in motor and dorsal premotor cortex of monkeys engaged in a delayed vibro-tactile discrimination task. Our findings are also somewhat reminiscent of results obtained in rat gustatory cortex in very different contexts. Specifically, Moran and Katz (2014) found that the transition among 2 HMM states evoked by sucrose were delayed by conditional taste aversion, while Mazzucato et al. (2019) found a faster onset of certain hidden states (named “coding states”) when a stimulus was expected, compared to the case where it was not expected. We will say more on this later.

### Slowdown of metastable dynamics in error trials

The other main result of this study is the link between a modulation of metastable activity and behavior. Specifically, we found that an increased duration of hidden states during error trials compared to correct trials. Moreover, this occurred during the deliberation period and not after the GO signal prompting the behavioral response. This result indicates that, during errors, one critical aspect that can be affected is the passage through states that presumably characterize different phases of the task, and before the action takes place. We are confident that the affected period reflects an internal deliberation because in our task the targets’ positions are revealed only at the GO signal, separating in time the deliberation phase from target selection and movement preparation.

Similarly, we found that reaction times were related to performance (correct vs. error) and not “objective” trial difficulty (where the latter is defined based on the relative distance between the two stimuli). It seems therefore that also the reaction times reflected the internal process of deliberation and whether such process resulted in correct performance, regardless of objective trial difficulty. These findings are reminiscent of intriguing results obtained in orbitofrontal cortex by Rich and Wallis (2016), who showed that slow deliberation was related to equal times spent in the latent states representing the available targets during a choice, rather than the difficulty of the decision as judged by eye movements. The closest analogy, however, is with the study of Mazzucato et al. (2019), who found shorter state durations when a stimulus was expected, compared to the case where it was not expected. The comparison with Mazzucato et al. (2019) invites speculation that correct decisions are more likely in trials in which expectations are successfully formed because, based on that study, this would predict faster dynamics in correct trials compared to error trials, as found here. In (Mazzucato et al., 2019) faster neural dynamics was related to faster stimulus decoding, not behavioral performance. The link with behavior was provided via a spiking network model, which showed that distractor stimuli could induce the opposite effect of slowing down the dynamics. If distractors would be more likely to induce an error, this would again predict slower dynamics prior to errors. Here we provide direct experimental evidence of a slowdown of the cortical dynamics during errors. To our knowledge, this is the first demonstration of a link between behavioral performance and the timescale of metastable dynamics in PFdl during a decision.

### Comparison with previous studies on error related activity in prefrontal cortex

The results reported here were obtained via an HMM analysis of ensemble activity. HMM analysis and its variations (Florian et al., 2011; Chen, 2013) present an ideal method to investigate the role of metastable dynamics. HMM allows an unsupervised segmentation of the ensemble activity without relying on external triggers, providing a characterization of the metastable dynamics during ongoing activity or motor preparation, not just following a stimulus (Abeles et al., 1995; Seidemann et al., 1996; Mazzucato et al., 2015). In particular, this allows one to pinpoint the moment in time in which an internal deliberation may have occurred. This is essential in trying to link behavior to neural activity occurring during internal deliberation as done here, and has allowed us to find neural correlates of trial difficulty and behavioral performance that had not emerged from single neuron analyses (Genovesio et al., 2011).

We note that our approach is very different from previous HMM studies of error activity, wherein an HMM fitted to trials in a given condition (say, S1>S2) was used to predict errors in a separate condition (S2<S1) (Ponce-Alvarez et al., 2012; Seidemann et al., 1996; Jones et al., 2007). In contrast, our ability to predict errors is based on the modulation of the dynamics of the same HMM. Our analysis on mean state durations uncovered a change in the network dynamics during incorrect trials that manifests itself as a reduced transition rate among successive states. Thus, errors are not predicted by the presence of specific “error states”, but rather by the slowing down of state sequences which may be the same in either condition.

Our approach is also different from previous studies which have examined PFdl activity during errors based on the activity of single neurons. From these studies the picture has emerged that PFdl neurons encode the adopted course of action during errors rather than the correct course of action. PFdl activity reflected the target chosen during errors in a motion discrimination task (Kim and Shadlen, 1999) and in a match-to-sample task for visual motion (Zaksas and Pasternak, 2006). Similar results were observed in tasks that required learning of action sequences, wherein PFdl neuronal activity reflected either the incorrect oculomotor sequence of actions (Averbeck and Lee, 2007) or the incorrect category of turn, pull, push hand movements during errors (Shima et al., 2007). At a more abstract level, PFdl neurons appear to encode goal and strategy in a task in which the goal was chosen after the selection of either a repeat-stay or a change-shift strategy (Genovesio et al., 2008). These strategies required to select the same or a different goal when a central instruction stimulus repeated or changed from the previous trial, respectively. PFdl neurons encoding the future goal appeared to encode the chosen goal rather than the correct goal during the decision period; however, strategy-coding neurons reflected the strategy adopted (rather than the correct strategy) only after the action. In Tsujimoto et al. (2011) a simplified version of the strategy task was adopted in which the shape and color of a cue stimulus indicated which strategy to use. This task required a more immediate strategy selection that was not based on the integration of previous events as in (Genovesio et al., 2005, 2008). In this case, during the decision period PFdl neurons encoded the strategy selected and not the strategy that had been cued. Interestingly, orbitofrontal neurons in the same task had the opposite behavior.

Finally, previous studies have shown the effect of previous trials on the activity of single neurons in the current trial. For example, Donahue et al. (2013) had shown the impact of reward on encoding of previous choice in multiple cortical areas, while a recent study by Spitmaan et al. (2020) has shown that both reward and choice outcomes are integrated over multiple trials to influence behavior and the response of cortical neurons. In the task studied here, we had found (Genovesio et al., 2014) that past outcome affected the reaction times but not the error rates, and only a very small effect of the previous choice on reaction times. These results suggest that, in our task, past outcome could have an influence on the metastable activity of PFdl neurons in the current trial. This influence could be revealed by the presence of coding states for the previous outcome. However, although single neurons encode both the previous choice and the previous outcome in the earlier part of the current trial prior to S2 (Genovesio et al., 2014), we found no evidence of coding states for previous outcome or choice during the temporal window analyzed in this paper (400 ms prior to S2 until the end of the trial; not shown). This suggests that the slowdown of dynamics between S2 and GO reflects the task demands of the current trial in a larger measure rather than the properties of the previous trial, although it cannot be ruled out that history effects could emerge in an HMM analysis performed in an earlier portion of the trial, or in datasets containing larger numbers of neurons.

### Link between single neurons and metastable ensemble activity

The studies reviewed above focused on the activity of single neurons taken individually rather than on ensemble dynamics as done here. It is natural to assume a link between the activity of single neurons and the sequence of states of the ensemble activity. We propose here one possibility based on our previous work on the evolution of goal representation in this task (Marcos et al., 2019). After the GO signal, we observed an abrupt reconfiguration of the prefrontal activity, in which the goal signal passed from one population to another as the trial proceeded from the delay period to the action period; only a smaller subset of neurons coded goals across both periods, and about half of these neurons switched goal preference between trial periods (“switch neurons”). These results were modeled by a network of heterogeneous cell assemblies with bistable local dynamics. In the model, the switch neurons showed a higher level of bursting activity than the other neurons, indicating that these neurons tended to jump more easily from high to low firing states, and vice versa. The switch neurons appeared also to have a leading role in the network reconfiguration after the GO signal, showing an earlier transition time in activity switch. It is possible that, when analyzed in terms of switching ensemble states via an HMM, the switch neurons could play an important role in igniting state transitions, and could act differently in correct vs error trials. The important connection between single cell and ensemble dynamics is left for future studies.

## Methods

In this study we used two adult, male rhesus monkeys (Macaca mulatta). Monkey 1 weighed 8.5 kg, and monkey 2 weighed 8.0 kg. All procedures followed the National Institutes of Health Guide for the Care and Use of Laboratory Animals (1996) and were approved by the National Institute of Mental Health Animal Care and Use Committee.

### Behavioral Task

In each trial, two visual stimuli were presented in sequence on a computer screen, separated by a temporal delay (Fig. 1a). Each stimulus could be either a blue circle of 3° diameter or a red square of 3 × 3° dimension. If the first stimulus (S1) was the red square then the second stimulus (S2) was the blue circle and vice versa. Each stimulus remained on screen for 1000 ms. Each trial started when the monkeys pressed the central of three switches, which caused the appearance on the screen of a central stimulus (reference point). After 400 or 800 ms the central stimulus was followed by the onset of S1. S1 was always located at a distance of 8–48 mm (in steps of 8 mm) above or below the reference point. After the disappearance of S1, there was a first delay (D1) of 400 or 800 ms before the presentation of S2. S2 appeared above the reference point if S1 had appeared below, and below the reference point otherwise. Its distance was as for S1, i.e., 8–48 mm (in steps of 8 mm) above or below the reference point, but it never equaled the distance of S1. Both S1 and S2 were presented for 1000 ms. The disappearance of S2 was followed by a second delay (D2) of 0, 400 or 800 ms, which in turn preceded the reappearance of the two stimuli, which served as a “GO” signal. Each stimulus was pseudo-randomly chosen to be located either 40 mm to the right or 40 mm to the left of the central stimulus. The monkeys were required to select, within 6 s, the stimulus presented farther from the reference point. Note that, not knowing the location of the stimuli, the monkeys could not plan any motor response before the GO signal. Correct choices were rewarded with 0.1 ml fluid, whereas an acoustic feedback followed the errors. All task variables, such as the duration of D1 and D2, the color and the shape of the two stimuli, were pseudorandomly determined. For more details about the behavioural task see Genovesio et al. (2011).

### Data collection

We monitored and recorded eye position with an infrared oculometer (Arrington Recording, Scottsdale, AZ, USA) and recorded single cells using quartz insulated, platinum-iridium electrodes (0.5–1.5 MΩ at 1 kHz), positioned by a 16-electrode drive assembly (Thomas Recording, Giessen, Germany). The electrodes were arranged within a concentric head with 518 mm spacing. Spikes were discriminated online using the Multichannel Acquisition Processor (Plexon, Dallas, TX, USA) and confirmed with the Offline Sorter (Plexon) based on principal component analysis, minimal interspike intervals, and well differentiated waveforms inspected individually for each isolated neuron.

### Surgery

We implanted the recording chambers over the exposed dura mater of the left frontal lobe, along with head restraint devices. We used Aseptic techniques together with isofluorane anesthesia (1–3%, to effect). Monkey 1 had two, 18 mm diameter chambers, and Monkey 2 had a single, 27×36 mm chamber.

### Histological analysis

Before the end of the recordings, electrolytic lesions (15 mA for 10 s, anodal current) were made at selected locations. After 10 days, the animal was deeply anesthetized and then perfused through the heart with formaldehyde-containing fixative (10% Formalin in 0.9% saline). We plotted recording sites on Nissl-stained coronal sections by reference to the recovered electrolytic lesions and the marking pins inserted during perfusion, and structural magnetic resonance images taken at various stages after the beginning of the recordings. Recordings were predominantly taken from area 8, area 46 and a small population of area 12.

### HMM analysis

An HMM was used to study the dynamics of neural ensembles, with methodology similar to (Ponce-Alvarez et al., 2012; Mazzucato et al., 2015, 2016, 2019). The sessions included in the analysis comprised at least 4 simultaneously recorded neurons, each with a mean trial activity ≥ 1 spk/s, and with at least 5 completed trials of each type, i.e., correct and incorrect. These selection rules left about 20% of the sessions for further analysis, out of the 361 initially recorded. Since many PFC neurons code for stimulus position (Genovesio et al., 2011), trials of the same session were divided according to the up or down position of the second stimulus, and were analyzed by fitting separate HMMs (in the analysis of trial difficulty of Fig. 6, similar results were obtained when fitting the same HMM to all correct trials in each session, see “Analysis of neural dynamics vs. trial difficulty”). In addition, in the analysis of correct vs. error trials (Figs. 2–5), HMMs were also fitted separately to correct vs. error trials in each session (see “Analysis of neural dynamics in correct vs error trials” below). In all HMM analyses described below, the vector of the neurons’ average firing rates across trials was used to initialize the emission rates for the first state. The same vector was then randomly permuted and assigned to the next state, until all states’ emission rates were initialized. In addition to a random permutation, a random Gaussian component with zero mean and 0.02 std was added to each emission rate. The transition matrix *P*_*ij*_, expressing the transition rates from state *i* to state *j* in each 5 ms bin, was initialized to 1 for the diagonal entries and |0.02*x*| off the diagonal, where *x* is a standard random variable; the rows of the matrix were then normalized to obtain probabilities. This initialization corresponds to a model where the probability is much larger to remain in the current state than to make a transition to another state in the next time bin. The fitting procedure was repeated 5 times with a maximum number of 500 iterations. The model with the smallest Bayesian Information Criterion (BIC) score, BIC = −2 LL + [*M* (*M* − 1) + *MN*] ln *T*, was selected as the model for further analysis, where LL is the log-likelihood of the model given the data, *M* is the number of hidden states, *N* is the number of neurons in the ensemble, and *T* is the number of observations in each session (number of trials × number of 5 ms bins per trial). A cross-validation procedure as performed e.g. in (Maboudi et al., 2018) gave similar results (not shown).

The training phase of the model, consisting in the estimation of the transition and emission probability matrices, was performed using the Baum-Welch algorithm. The states were then decoded from the posterior state probabilities *P* (*S*_*t*_|*X*_*t*_) of having state *S*_*t*_ in the presence of data *X*_*t*_ (spike trains) in a 5 ms time bin centered around time *t*. We assigned a state to a chunk of data only if its posterior probability was greater than 0.8 for at least 50 consecutive ms, as this requirement reduces the chance of overfitting the data (Mazzucato et al., 2015, 2016).

All analyses were performed using custom software written in Python and Matlab (Statistics and Machine Learning Toolbox, ©The Mathworks). The code is available at *https://github.com/danilobenozzo/hmm_neurofis.git*

### Coding states

The presence of a coding state, meaning a state that codes for a specific task condition, was evaluated by a *χ*-squared test of the frequency of appearance of each state in a given task condition during a time window of interest. The frequency table was computed by reporting the frequency of occurrence of each state in the two task conditions to be compared. As task condition, we considered the relative distance of the two stimuli from the center based on stimulus features (blue circle vs. red square) and order of presentation (S2 farther vs. closer), and position of the target at GO (right vs. left; the target is the stimulus previously presented farther from the center). As for the time window of interest, we chose the S2 presentation period for states coding for relative distance based on stimulus features or order, and the time interval from GO to RT for the target position.

### Shuffled datasets

The HMM analysis was validated via comparison with shuffled data. Three types of shuffling procedures were applied in each session: a circular shuffle, a swap shuffle, and class label shuffle. In the latter, the class labels associated with each trial type (S1>S2 vs. S2>S1) were randomly shuffled. Circular shuffle (Maboudi et al., 2018) consisted of independent random time shifts of the spike trains in each trial. This procedure kept the single neuron autocorrelations intact but disrupted the cross-correlations across neurons and, in particular, instances of co-activation. Swap shuffles (Maboudi et al., 2018) randomly permuted the temporal bins, i.e., the vectors of spike counts, in each trial. This procedure preserved the neurons’ cross-correlations but removed the neurons’ autocorrelations, in addition to the order of the sequential patterns that might be present in the data. Note that neither procedure changed the overall firing rates of the neurons in each trial. The HMM fits of the shuffled datasets were compared with the fit of the original dataset in terms of BIC score and optimal number of states. For the sake of visualization, in Fig.3a-b-c time has been uniformly warped in the main task epochs (e.g. S2 presentation, delay D2, reaction time) by a normalisation with respect to each epoch length in order to align the related events across trials.

### Analysis of neural dynamics in correct vs error trials

To compare state durations in correct and error trials, the HMM was fitted to all data inside a time window of variable size starting 400 ms before S2 and ending at the beginning of the following trial. Trials of the same session were divided according to the up or down position of the second stimulus and into correct vs. error, and were analyzed by fitting four separate HMMs in each condition (56 sessions in total). In each session, we used models with 2 to *N* − 1 states, where *N* is the number of neurons in the session (the number of states was kept lower than the number of neurons to help prevent overfitting). The optimal number of states in each session, *M*, was chosen so as to minimize the BIC score (see formula reported above). Two separate HMMs were fitted to correct and incorrect trials in the same session given the stimulus position. After decoding the data according to the posterior state probabilities *P* (*S*_*t*_|*X*_*t*_) as described above, we compared the session-averaged state durations in correct vs. error trials in two different time epochs, from offset of S2 to GO, and from GO to END, by performing a 2-way ANOVA with factors trial type (correct vs. error) and time epoch. To validate the analysis with a balanced number of trials in each trial type (correct vs. error), we randomly selected a number of correct trials matching the number of error trials in each session. The HMM analysis was performed on the trial-matched sessions as described above, and the mean state durations in correct vs. error trials across all sessions were obtained. This procedure was repeated 20 times, each time with a new pseudo-random selection of correct trials. These same trial-balanced datasets were then analyzed after circularly shuffling each spike train (see Section “Shuffled datasets”). Swap-shuffling the data drastically reduced the number of states and transitions and resulted in only a handful of trials with at least 2 transitions in each session, precluding a meaningful analysis of the state durations in this case.

### ROC analysis

We used the distributions of mean state durations from S2 to GO obtained as described in Section “Analysis of neural dynamics in correct vs error trials” to predict the performance (correct vs. error) in single trials. To do so, we treated the task of predicting performance as a fictitious two-alternative forced choice (2AFC) task in which we are presented with a correct and an error trial, and for each such pair, we predict that the error trial is the one with longer mean state duration. Performance in this task can be obtained by computing the area under the receiver operating characteristic (ROC) curve (Dayan and Abbott, 2005). The latter is the curve plotting the fraction of true positives (hits) vs. the fraction of false positives (false alarms) obtained as the decision threshold varies from +∞ to −∞, where the task is to classify single trials and each decision is based on whether the trial’s mean state duration is above or below the threshold. Note that this analysis is analogous to that performed in (Britten et al., 1992) for decoding motion direction from single neurons’ firing rates. However, in (Britten et al., 1992) the authors aimed to decode the stimulus while we aimed to decode the monkeys’ performance (correct vs. error). In the latter case, the prediction of a correct trial corresponds to matching the monkeys’ behavior in that trial. Thus, our average decoding performance also gives the probability of predicting the monkey’s behavior on a trial-by-trial basis.

### Analysis of neural dynamics vs. trial difficulty

For this analysis, we fitted an HMM to the data inside a 1400 ms time window (400 ms before the presentation of S2 until its removal), and the number of states was set to 2 (Ponce-Alvarez et al., 2012). Only correct trials were used, divided according to the up or down position of the second stimulus, and then analyzed by fitting separate HMMs (however, similar results were obtained when fitting the same HMM to all correct trials in each session; not shown). We defined the time until the first transition after S2 (fTaS2) as the period of time between S2 onset and the time at which the posterior probability of the current state decreases below 0.8 (see Fig.6 for examples). fTaS2 times were regressed with |S2-S1|as described in Results. In this analysis, only sessions with significant decoding performance by the HMM were considered (41 out of 61 sessions with S2 up and 34/61 sessions with S2 down), according to the following procedure (Ponce-Alvarez et al., 2012). We divided each session in testing and training sets in a 3-fold cross-validation framework. For each trial in the testing set, its log-likelihood was computed both by a model fitted in the training trials of class S1>S2 (model *λ*_1_) and by a model fitted in the training trials of class S2>S1 (model *λ*_2_; here, “S1>S2” means that S1 was more distant than S2 from the central stimulus). The tested trial was assigned to class S1>S2 if model *λ*_1_ had the larger log-likelihood on this trial, and to class S2>S1 otherwise. The session was kept for further analysis if the decoding performance of the above method, measured by balanced accuracy, was significantly larger than chance. Balanced accuracy was defined as the average of recall obtained on each class, i.e. 1*/*2(TP/P+TN/N), and was used because in most sessions the two classes S1>S2 and S2>S1 were unbalanced (i.e., there were an unequal number of trials in each class; see (Brodersen et al., 2010) for details). Here, TP means “true positives”, TN means “true negatives”, and P, N stands for positive (i.e., elements of class S1>S2) and negative (elements of class S2>S1), respectively. Significance of the decoder’s balanced accuracy was computed by a binomial test at the 5% significance level. The same analysis was performed after shuffling the data as described in Section “Shuffled datasets”.

## Acknowledgements

We thank Dr. Steven P. Wise, Dr. Satoshi Tsujimoto and Andrew Mitz for their contribution in the experimental phase of the study. This work has been partially supported by a grant from the Sapienza University (Progetto H2020: PH1181642DB714F6) to A.G. and by an NIH/NINDS Brain Initiative grant (1UF1NS115779) and a grant from the Office of the Vice President for Research of Stony Brook University (award #63845) to G.L.C. The data analyzed in this article were collected at the National Institute of Mental Health with support from the Division of Intramural Research. The views expressed in this article do not necessarily represent the views of the NIMH, NIH, or the United States Government.

## Author contributions statement

A.G. designed the experimental task and collected the data. D.B., G.L.C and A.G. designed the data analysis and wrote the paper. D.B. performed research. G.L.C and A.G. supervised research.

